# Trophic consistency of supraspecific taxa in belowground invertebrate communities

**DOI:** 10.1101/455360

**Authors:** Anton M. Potapov, Stefan Scheu, Alexei V. Tiunov

## Abstract

1. Animals that have similar morphological traits are expected to share similar ecological niches. This statement applies to individual animals within a species and thus species often serve as the functional units in ecological studies. Species are further grouped into higher-ranked taxonomic units based on their morphological similarity and thus are also expected to be ecologically similar. On the other hand, theory predicts that strong competition between closely related species can lead to differentiation of ecological niches. Due to a high diversity and limited taxonomic expertise, soil food webs are often resolved using supraspecific taxa such as families, orders or even classes as functional units.

2. Here we for the first time empirically tested the trophic consistency of supraspecific taxa across major lineages of temperate forest soil invertebrates: Annelida, Chelicerata, Myriapoda, Crustacea and Hexapoda. Published data on stable isotope compositions of carbon and nitrogen were used to infer basal resources and trophic level, and explore the relationship between taxonomic and trophic dissimilarity of local populations.

3. Genera and families had normal and unimodal distributions of isotope niches, suggesting that supraspecific taxa are trophically consistent. The isotopic niche of populations across different localities is better predicted by species than by supraspecific taxa. However, within the same genus, the effect of species identity on stable isotope composition of populations was not significant in 92% of cases. The link to basal resources, i.e. plants or detritus, was convergent in different lineages, while trophic levels followed the Brownian motion taxonomic model. Virtually none of the studied taxa showed pronounced trophic niche conservatism within a lineage.

4. Supraspecific taxa are meaningful as functional units in ecological studies, but the consistency varies among taxa and thus the choice of taxonomic resolution depends on the research question; generally, identification of taxa should be more detailed in more diverse taxonomic groups. We compiled a comprehensive list of mean Δ^13^C and Δ^15^N values of invertebrate taxa from temperate forest soils allowing to refine soil food-web models when identification to species level is not feasible.

## Introduction

Shared ancestry typically is associated with ecological similarity, with closely related species having similar traits and therefore similar ecological niches (Webb, 2000; Webb, Ackerly, McPeek, & Donoghue, 2002). Such similarity results from ‘evolutionary inertia’ and is referred to as phylogenetic signal (Blomberg & Garland, 2002). The inertia is likely supported by the ancestral constraint mechanisms, i.e. the property of a trait that, although possibly adaptive in the environment in which it originally evolved, acts to place limits on the evolution of new phenotypic variants (Blomberg & Garland, 2002; Pyron, Costa, Patten, & Burbrink, 2015). Notably, not only the trait itself can be conserved but also its potential to vary. This concept was formalized in the early 20th century by N. Vavilov in “*The law of homology series in genetical mutability”* (1935) and implies that related species tend to evolve similar phenotypic traits.

As a result of evolutionary inertia, taxonomically related species may respond similarly to environmental factors and perform similar ecosystem functions, since the organisms that share similar morphology are likely to share a similar ecological niche (Wainwright & Richard, 1995). Moreover, supraspecific taxa are suggested to behave as ecological units forming the structural parts of ecosystems (Chernov, 2008). The taxonomic level to which ecological characteristics of species can be extrapolated with a minimum loss of information, i.e. “taxonomic sufficiency”, is an important question in basic and applied ecology (Ellis, 1985; Terlizzi, Bevilacqua, Fraschetti, & Boero, 2003). For instance, a moderate level of taxonomic aggregation (genera or even families) can be sufficient to indicate effects of environmental changes on the composition of benthic and soil invertebrates communities (Jiang et al., 2013; Minor, Ermilov, & Tiunov, 2017). Loss of information due to taxonomic aggregation likely is more pronounced in more diverse groups (Timms, Bowden, Summerville, & Buddle, 2013), but the ecological consistency of taxa at different taxonomic levels has never been tested empirically.

Ecological similarity among related species usually is investigated in respect to morphology, physiology and behaviour of organisms, with behavioural adaptations being evolutionarily more labile (Blomberg, Garland, Ives, & Crespi, 2003; Böhning-Gaese & Oberrath, 1999). As a complex trait, trophic niches may either be similar or differ considerably among closely related taxa. Feeding habits of an organism are not strongly related to its taxonomic position since similar feeding strategies occur in many lineages (i.e., convergence in trophic positions). Pronounced trophic-niche differentiation between species was shown for dominant groups of soil detritivores (Chahartaghi, Langel, Scheu, & Ruess, 2005; Schneider et al., 2004). Nevertheless, supraspecific taxa often are used as trophic species (nodes) in soil food web models (Berg & Bengtsson, 2007; de Ruiter, Neutel, & Moore, 1994; de Vries et al., 2013). Despite trophic guilds conceptually are independent of taxonomy, taxonomically related species usually are included in the same trophic guild (Simberloff & Dayan, 1991). This assumption recently has received empirical support in freshwater (catfishes) and soil (springtails) detritivores, for which trophic niches were shown to differ between supraspecific taxa (Lujan, Winemiller, & Armbruster, 2012; Potapov, Semenina, Korotkevich, Kuznetsova, & Tiunov, 2016). Moreover, it has been argued that ecological links between species (e.g., via trophic interactions) often are related to their phylogenetic position (Gómez, Verdú, & Perfectti, 2010; Rezende, Lavabre, Guimarães, Jordano, & Bascompte, 2007). In marine communities predators tend to consume phylogenetically related prey species and prey tend to be consumed by phylogenetically related predators (Eklöf & Stouffer, 2016). However, it remains unclear if this also holds true for taxa in terrestrial belowground communities with less pronounced size structure and high frequency of omnivorous species able to switch diet depending on its availability (Digel, Curtsdotter, Riede, Klarner, & Brose, 2014).

Three scenarios are suggested to explain the relationship between taxonomic relatedness of taxa and their dissimilarity in trophic niche. The ‘limiting similarity’ hypothesis suggests that morphological similarity of closely related taxa results in strong competition (Darwin, 1876; Violle, Nemergut, Pu, & Jiang, 2011) and this may favour trophic niche divergence. Although trophic niches of species may be dissimilar, shared ancestral morphology should lead to the overlap of trophic niches of supraspecific taxa, such as genera and families (Fig. 1). The ‘random’ hypothesis suggests no changes in trophic-niche differentiation at different levels of taxonomic resolution (i.e., morphological dissimilarity is not related to trophic niche dissimilarity). The ‘taxonomic signal’ hypothesis suggests that closely related taxa have more similar trophic niches than distantly related taxa. Accordingly, supraspecific taxa are expected to occupy distinct trophic niches that reflect ancestral adaptations, while genera within families and species within genera differ only little.

**Figure 1.**
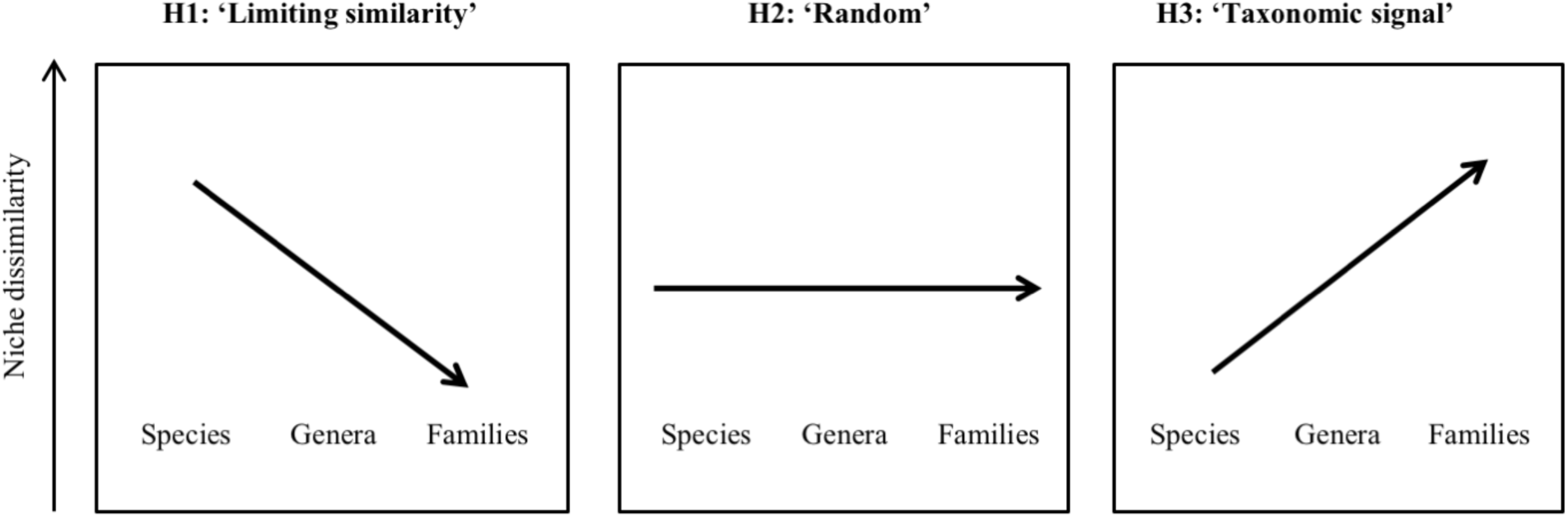
Contrasting scenarios of niche differentiation among closely-related taxa. Hypothesis 1 (‘limiting similarity’) suggests strong competition and trophic-niche divergence among closely-related taxa. Hypothesis 2 (‘random’) hypothesis suggests no relationship between taxonomic and trophic-niche dissimilarity. Hypothesis 3 (‘taxonomic signal’) suggests that closely related taxa have more similar trophic niches than distantly related taxa and that niche differentiation is increasing with increasing taxonomic level.

The limitations of measuring trophic-niche dimensions *in situ* were partly overcome with the introduction of stable isotope analysis in ecological studies (Peterson & Fry, 1987). Natural variations in stable isotope ratios of carbon (^13^C/^12^C) and nitrogen (^15^N/^14^/N) reflect the trophic niche of species (Newsome, Martinez del Rio, Bearhop, & Phillips, 2007; Rodríguez & Herrera, 2013) and provide information on their basal resources and trophic level, respectively (Boecklen, Yarnes, Cook, & James, 2011; Martinez del Rio, Wolf, Carleton, & Gannes, 2009). In soil food webs ^13^C concentrations increase with decomposition of plant material, while ^15^N concentrations increase with trophic level (Potapov et al., 2018). Based on these findings, we used published data on stable isotope composition of carbon and nitrogen to explore the trophic niches of major belowground arthropod lineages at different taxonomic levels. The following main questions were addressed: (1) Is there evidence for trophic niche conservatism (*sensu* Losos, 2008) in soil animal taxa? (2) Are supraspecific taxa in soil communities trophically consistent? (3) How much information about basal resources and trophic level of soil animals do we lose by using supraspecific taxa instead of species?

## Methods

The analysis was based on a large compilation of published data on stable isotope composition of soil invertebrates from temperate forests (Potapov et al., 2018). In the dataset, only records identified to species or genus level were included resulting in 415 species from 21 orders and 5 high-order taxa: Annelida, Chelicerata, Myriapoda, Crustacea and Hexapoda (Table S1). Each data record in the dataset comprised average stable isotope composition of species / taxa sampled at a local community with some species investigated at different localities / ecosystems resulting in a total of 961 local populations. Stable isotope compositions of carbon and nitrogen were normalized to the local leaf litter to account for inter-ecosystem variations and denoted as Δ^13^C and Δ^15^N values (Klarner et al., 2014). Further, each record was classified into the following taxonomic levels: Phylum, Subphylum, Class, Order, Family, Genus and Species, according to (Zhang, 2011). Although arbitrarily defined and varying between taxonomic groups, supraspecific taxa form a hierarchical system representing different levels of aggregation of species.

Statistical analyses were performed in R 3.4.0 with R studio interface (R Core Team, 2017). To test if stable isotope composition of soil invertebrates is distributed non-randomly on the taxonomic tree, we analysed Δ^13^C and Δ^15^N values using Blomberg’s K criterion in the package *phytools* (Revell, 2012). This criterion was developed to test for significance of phylogenetic signal, i.e. its non-random distribution on a phylogenetic tree, and allows comparisons of phylogenetic signal across traits and phylogenetic trees (Blomberg et al., 2003). Data were transformed to a tree using *as.phylo* in package *ape* (Paradis, Claude, & Strimmer, 2004). The empirical Blomberg’s K criteria were compared with 999 Brownian motion simulations (*fastBM*) in package *phytools*. Due to a positive skewness of the distribution, the 95% confidence interval for simulated K criteria was calculated after log-transformation and was estimated as 0.62-1.81. K values below 0.62 indicate trophic-niche convergence among taxa (similar isotopic composition is observed in different lineages), while values above 1.81 indicate trophic-niche conservatism (different lineages have different isotopic composition; Losos, 2008).

According to the abovementioned hypotheses, we explored the relationship between isotopic dissimilarity and taxonomic dissimilarity of taxa at different taxonomic levels. Isotopic dissimilarity (as a proxy of trophic niche dissimilarity) was measured as Euclidean distance between populations’ Δ^13^C and Δ^15^N values. Taxonomic dissimilarity was ranked from 1 to 8, with 1 denoting different populations of the same species, 2 denoting populations of different species of the same genus, 3 – populations of different genera of the same family, 4 – populations of different families of the same order, 5 – populations of different orders of the same class, 6 – populations of different classes of the same subphylum, 7 – populations of different subphyla of the same phylum, and 8 – populations of different phyla. Matrixes for isotopic and taxonomic dissimilarities were calculated using *daisy* in package *cluster* (Maechler, Rousseeuw, Struyf, Hubert, & Hornik, 2017). Subsequently, isotopic and taxonomic dissimilarity matrixes were correlated using Mantel test with 999 permutations (package *ade4*; (Dray, Dufour, & others, 2007). Medians and means of isotopic dissimilarity were compared between taxonomic levels using Nemenyi test in package *PMCMR* (Pohlert, 2014) and general linear hypothesis for multiple comparisons in package *multcomp* (Hothorn, Bretz, & Westfall, 2008), respectively. Both tests showed similar results; only results of the former are presented.

To test the trophic consistency of supraspecific taxa we visually explored the distributions of Δ^13^C or Δ^15^N values of all taxa at different taxonomic levels. To analyse all taxa together, we first calculated mean Δ^13^C and Δ^15^N values for all taxa at all taxonomic levels. Next, we calculated offsets of Δ^13^C and Δ^15^N values for populations from the mean values for all parental taxa. For example, to construct data distribution for the ‘family’ level, offsets for all populations were calculated using the mean values of the corresponding parental families. Only taxa with two or more nested taxa of the lower taxonomic level were included in the analysis (i.e., species with ≥ 2 populations, genera with ≥ 2 species, families with ≥ 2 genera etc.). Offsets were compared between taxonomic levels using general linear hypothesis for multiple comparisons in package *multcomp*.

To estimate how much information about basal resources and trophic level of soil animals is lost by using supraspecific taxa instead of species, we checked whether different species within the same parental taxon occupy significantly distinct trophic niches. For doing that, we inspected the effect of ‘species identity’ on the Δ^13^C or Δ^15^N values of populations using one-way ANOVAs. Only taxa with two or more nested taxa of the lower taxonomic level were included in the analysis. Thereafter, results of the ANOVA (effect of species / no effect of species as a binomial variable) were used to calculate percentage of taxa with significant species effect at different levels of taxonomic resolution.

Lastly, we inspected mean Δ^13^C and Δ^15^N values for individual subphyla, classes and orders across three major lineages of soil arthropods that were well represented in the dataset, i.e. Chelicerata, Myriapoda and Hexapoda. In this analysis we used the expanded dataset of 1146 populations including those not identified to species level (Table S1).

## Results

Based on the entire dataset both Δ^13^C and Δ^15^N values showed taxonomic signal, i.e. closely related taxonomic units shared basal resources and trophic levels more often than expected by random distribution (p = 0.001 for both). Comparisons based on the Brownian motion model showed that trophic niches are not taxonomically conserved with Δ^13^C values being convergent among taxa (Blomberg’s K = 0.50) and Δ^15^N values distributed following the Brownian motion model (K = 0.87).

Mean isotopic dissimilarity between populations in Δ^13^C values only slightly increased with taxonomic dissimilarity (Mantel test: R = 0.006, simulated p = 0.245); by contrast, isotopic dissimilarity in Δ^15^N values increased steadily from the population to the class level but decreased thereafter (R = 0.065, p = 0.001; Fig. 2). Whereas populations of different species within a genus differed by 1.3‰ (a median), populations from different classes within a subphylum differed by 3.9‰.

**Figure 2.**
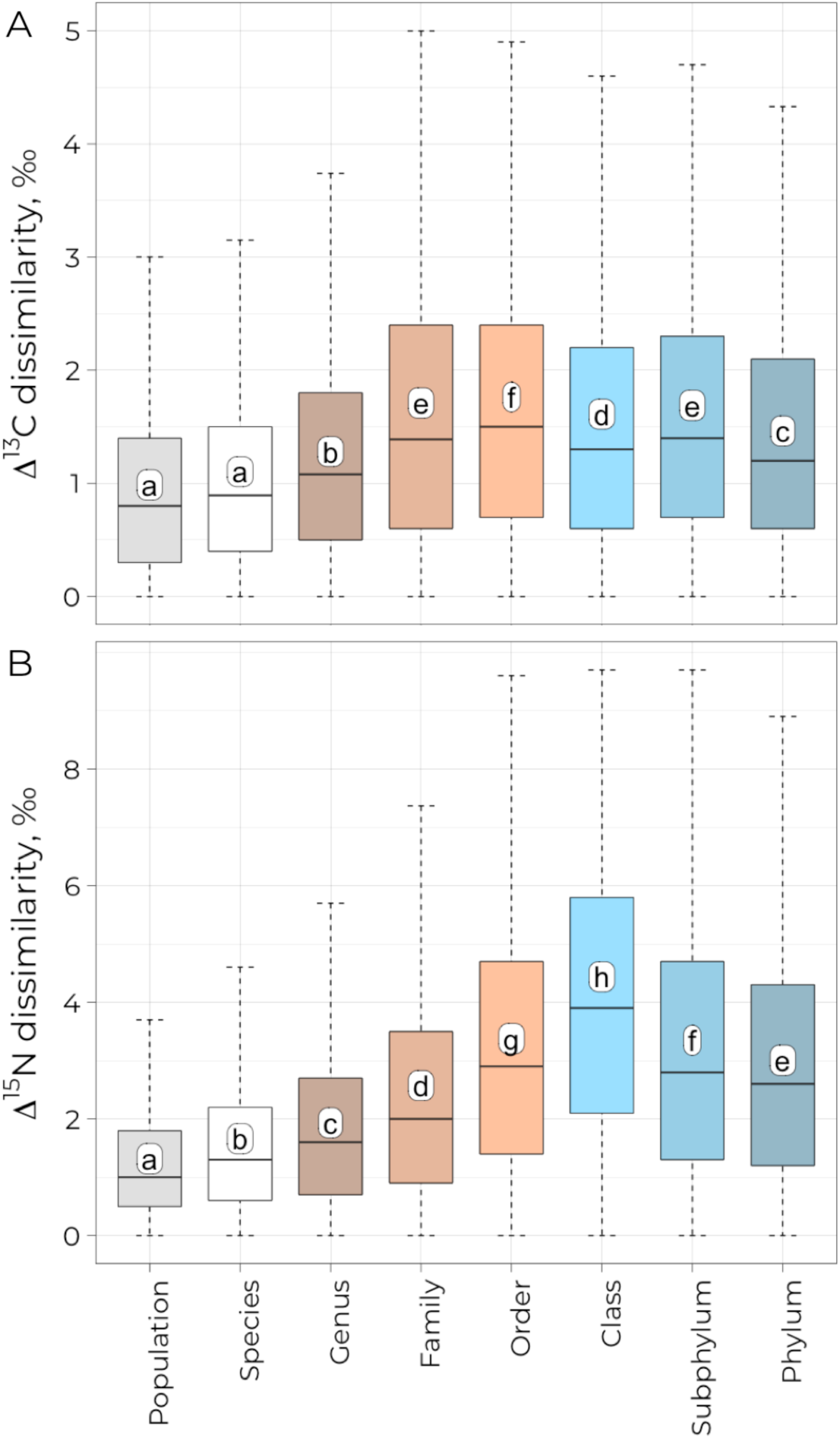
Isotopic dissimilarity as related to taxonomic dissimilarity in soil invertebrates. Isotopic dissimilarity was measured as Euclidean distance in Δ^13^C or Δ^15^N values between populations. Taxonomic dissimilarity is reflected by eight levels of taxonomic resolution: For instance, ‘Population’ represents differences between populations of the same species, ‘Species’ represents differences between populations from different species of the same genus, ‘Genus’ represents differences between populations from different genera of the same family, etc. Distances within groups are illustrated using boxplots with horizontal lines representing medians; long upper tails in few cases were cut for clarity (2% of records for both Δ^13^C and Δ^15^N). Means are indicated by white circles; letters refer to median comparison (Nemenyi test): groups sharing the same letter are not significantly different. Isotopic and taxonomic dissimilarity were significantly correlated for Δ^15^N but not for Δ^13^C values (see text).

The variability of both Δ^13^C and Δ^15^N values increased continuously with decreasing level of taxonomic resolution. However, taxa from species to family level for Δ^15^N and from species to order level for Δ^13^C fitted normal and unimodal distribution (Fig. 3A, B), suggesting trophic consistency of taxa at these taxonomic levels. The offset of individual populations from the mean values of parental taxa continuously increased with decreasing taxonomic resolution until class and subphylum levels for Δ^13^C and Δ^15^N values, respectively (Fig. 3C, D). Average absolute offsets of Δ^13^C values were 0.6‰ for species and 0.8‰ for families; average offsets of Δ^15^N values were 0.9‰ for species and 1.3‰ for families.

**Figure 3.**
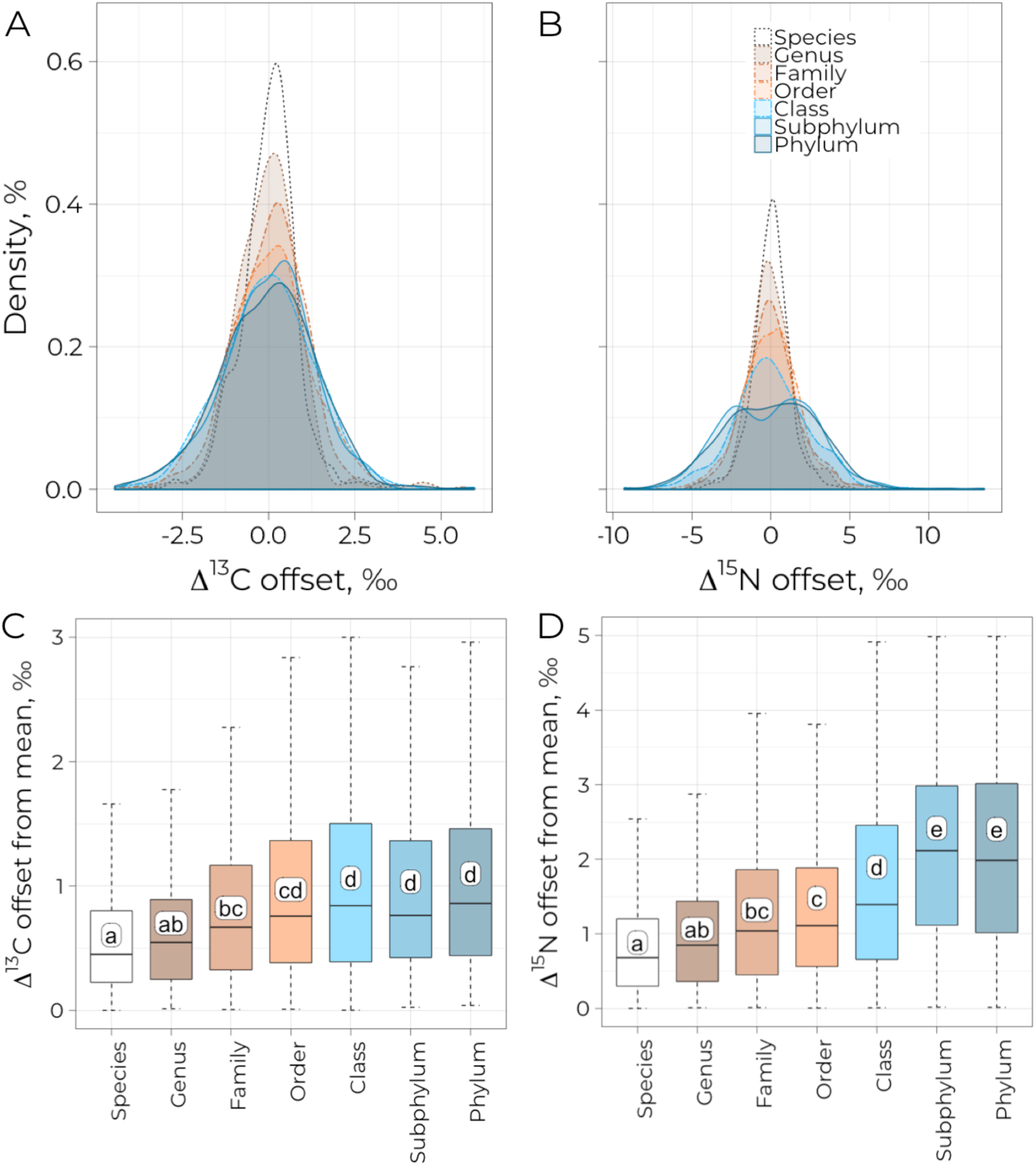
Offsets in stable isotope composition of populations from means of parental taxa of different taxonomic level. Offsets were calculated as mean Δ^13^C (A, C) and Δ^15^N values (B, D) for populations minus the mean value for parental taxonomic units, all data are plotted together. E.g. ‘Family’ stands for offsets of individual populations from the mean of parental families. Only taxa with two or more nested taxa of the lower taxonomic level were included in the analysis (i.e., species with ≥ 2 populations, genera with ≥ 2 species, families with ≥ 2 genera etc.). A and B: Offsets are displayed as the Kernel density estimation. C and D: Absolute offsets are displayer as boxplots with horizontal lines representing medians. Means are indicated by white circles; letters refer to median comparison (Nemenyi test): groups sharing the same letter are not significantly different.

To estimate, to what extent information is lost by using supraspecific taxa instead of species, we tested for the effect of species on Δ^13^C and Δ^15^N values within each supraspecific taxa from genus to class level. Within genera, the effect of species was not significant in 92% of the cases for both Δ^13^C and Δ^15^N values (Supplementary Table S2). Within families, orders, and classes the effect of species on Δ^13^C values was not significant in 82, 45, and 20% of cases, respectively. The respective values for Δ^15^N values were 59, 45, and 0% (Fig. 4).

**Figure 4.**
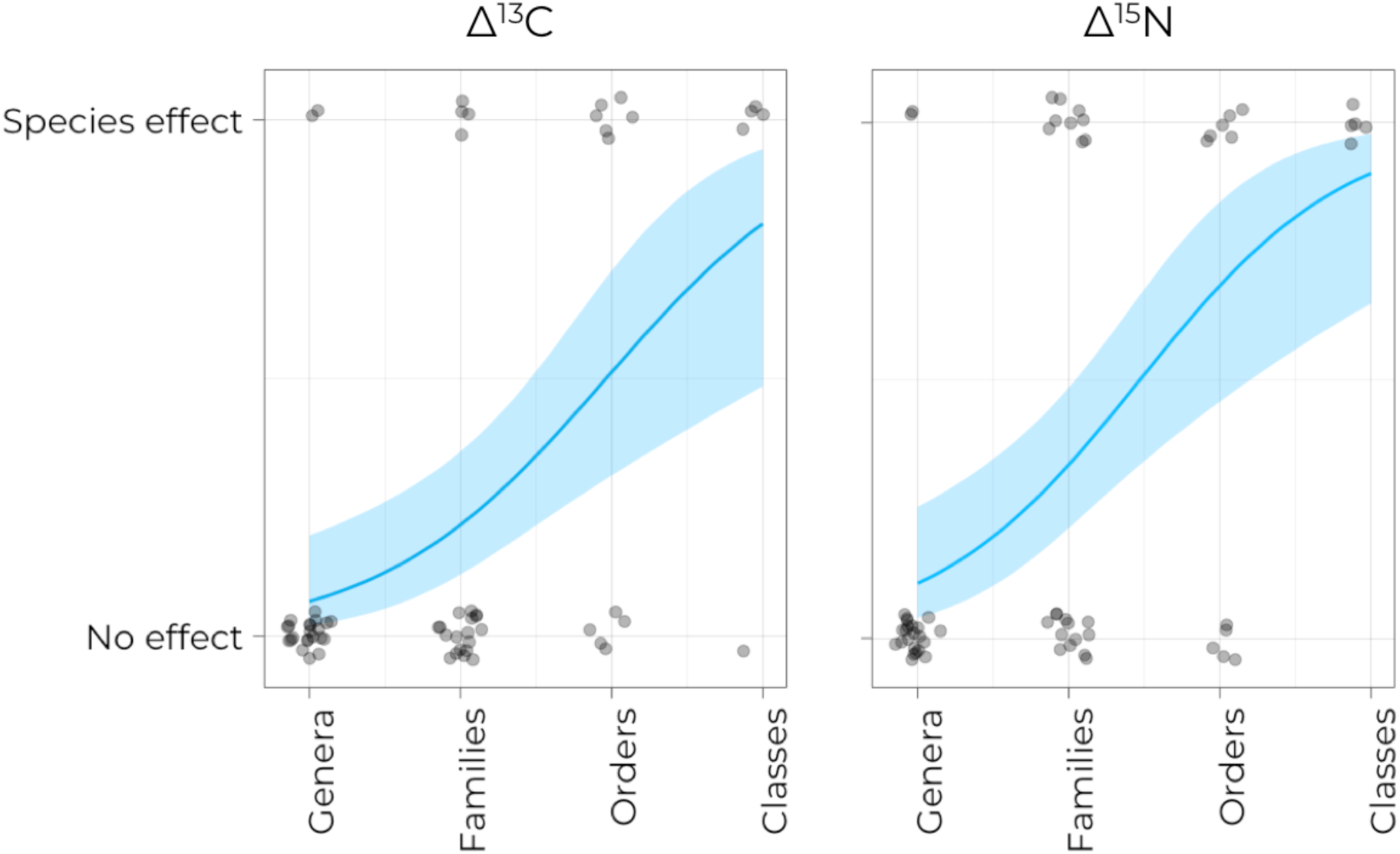
Effect of species identity on Δ^13^C and Δ^15^N values at different taxonomic levels. Significant effects of species on population stable isotope composition within the parental taxon were inspected using ANOVA (Table S3). Only taxa with two or more nested taxa of the lower taxonomic level were included in the analysis (i.e., species with ≥ 2 populations, genera with ≥ 2 species, families with ≥ 2 genera etc.). Probability of significant effect is illustrated with binomial smoother; shaded areas show 95% confidence intervals.

More detailed analysis of high-rank taxa showed that at the level of subphyla, Chelicerata and Myriapoda were on average enriched in both ^13^C and ^15^N as compared to Hexapoda (Figs 5, 6). There were no differences in Δ^13^C values between classes within subphyla, but classes within Myriapoda and Hexapoda were distinct in Δ^15^N values (Chilopoda vs. Diplopoda and Insecta vs. Collembola). Further, Δ^13^C values differed between orders within Arachnida, Insecta and Collembola, but not within Chilopoda and Diplopoda. Δ^15^N values clearly differed between predominantly predatory orders (Mesostigmata, Araneae, Geophilomorpha, Lithobiomorpha, Coleoptera and Hymenoptera) and predominantly detritivore orders (Oribatida, Julida, Polydesmida, Entomobryomorpha and Symphypleona). Diptera and Poduromorpha had intermediate Δ^15^N values. Significant differences were also found at the family and genus level of taxonomic resolution (Supplementary materials, Fig. S1-S3).

**Figure 5.**
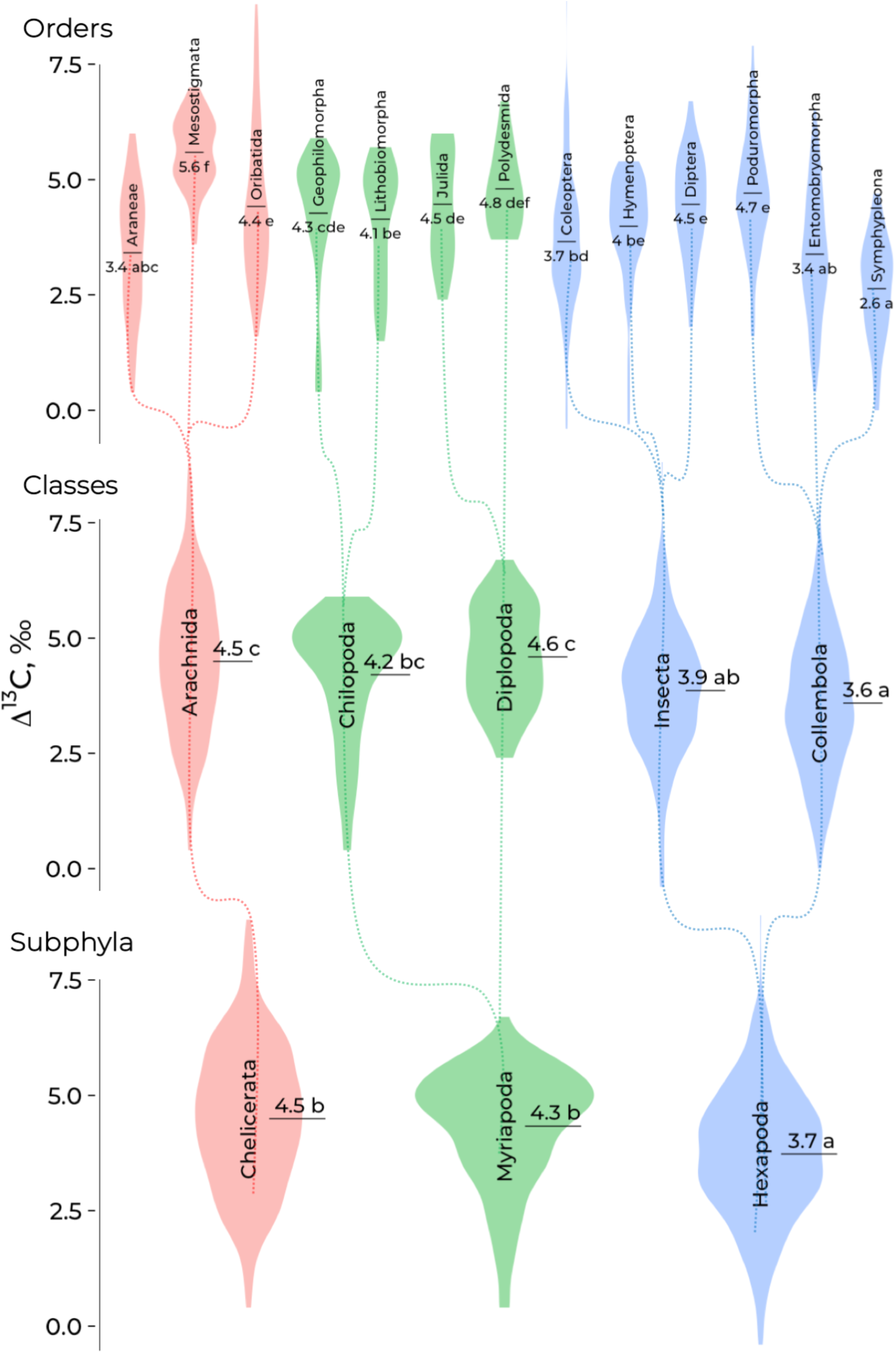
Δ^13^C values of subphyla, classes and orders in three main lineages of soil arthropods, Chelicerata, Myriapoda and Hexapoda. Data on populations are presented in the form of violin plots (mirrored Kernel density estimation). Mean values are shown with horizontal segments and numbers; mean values sharing the same letter within a taxonomic level are not significantly different (general linear hypothesis for multiple comparisons). Colours denote different lineages, dotted lines denote taxon hierarchy.

**Figure 6.**
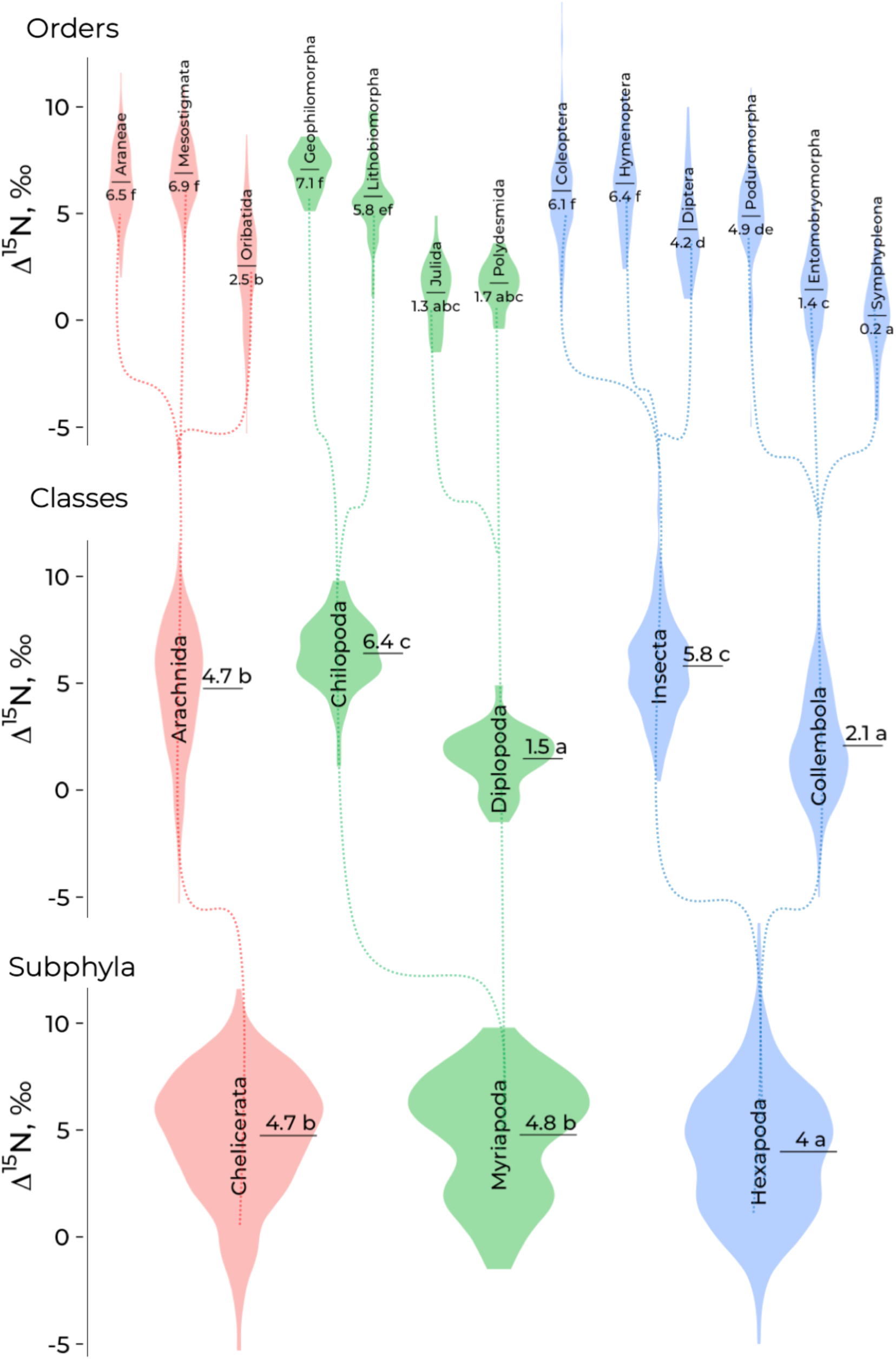
Δ^15^N values of subphyla, classes and orders in three main lineages of soil arthropods, Chelicerata, Myriapoda and Hexapoda. Data on populations are presented in the form of violin plots (mirrored Kernel density estimation). Mean values are shown with horizontal segments and numbers; mean values sharing the same letter within a taxonomic level are not significantly different (general linear hypothesis for multiple comparisons). Colours denote different lineages, dotted lines denote taxon hierarchy.

## Discussion

Supraspecific taxa are often assumed to comprise species that perform similar ecological function. However, this assumption has not been formally tested. Using stable isotope analysis we found strong support that dissimilarity in trophic level in soil invertebrates is related to taxonomic dissimilarity, supporting the ‘taxonomic signal’ hypothesis. Trophic level (as indicated by Δ^15^N values) was well reflected by taxonomic units, which was mainly due to the fact that high-order taxa (i.e., class and order level) encompass either detritivores or predators. The pattern in Δ^13^C values supported ‘random’ hypothesis, indicating that similar basal resources can be utilized by various taxonomic groups with different types of body organisation.

The distribution of Δ^13^C and Δ^15^N values in supraspecific taxa, at least for genera and families, were on average unimodal and close to normal, indicating that supraspecific taxa are trophically consistent. Depending on the research question, our finding validates the usage of supraspecific taxa as functional units in ecological studies if identification to species level is not feasible. As expected low-rank taxonomic units provide a higher precision and a lower chance of information loss. Still, due to a high variability within species, the effect of species identity on Δ^13^C and Δ^15^N values of populations was not significant in 92% of the tested genera.

### Basal-resource and trophic-level conservatism in soil food webs

Since we analysed a taxonomic rather than a phylogenetic tree, our conclusions about evolutionary aspects of trophic niche conservatism are limited. However, the taxonomic tree that we used reflects well the basic distinction between the main invertebrate lineages/high-rank taxa (Misof et al., 2014; Rehm et al., 2011; Rota-Stabelli, Daley, & Pisani, 2013). Thus, the prominent differences in trophic niches for high-rank taxa (orders, classes) suggest that early evolutionary adaptations played a crucial role in shaping the niches of soil invertebrates. Similarly, in arbuscular mycorrhizal fungi different colonization and resource exploitation strategies evolved early during diversification (Powell et al., 2009).

Phylogenetic inertia constrain the evolution of ecological niches, but this can be very different for different niche dimensions. For instance, behavioural adaptations in vertebrates are quite labile, while morphological adaptations are often evolutionary conserved (Blomberg et al., 2003; Böhning-Gaese & Oberrath, 1999). The concept of evolutionary conservatism was refined by Losos (2008) who emphasized that ecological niches are often related to phylogeny and the question is whether they are more or less conserved than expected by the Brownian motion model. Both Δ^13^C and Δ^15^N values were distributed on the taxonomic tree in a non-random way, however, we found no evidence that trophic niches are more conserved than expected by Brownian motion. Δ^13^C values (reflecting the link to basal resources) were likely evolutionary convergent in different lineages. Δ^15^N values (reflecting the trophic level) followed the Brownian motion.

It has been suggested that niche conservatism is reinforced by stabilizing selection due to the presence of sympatric species which occupy adjacent ecological niches (Ackerly, 2003; Losos, 2008). Despite soil is one of the most densely populated habitats on Earth, trophic niche conservatism is not evident. Presumably, the heterogeneous nature of soil provides an array of microhabitats varying in space, size and time, and therefore reduces competitive interactions between species (Maaß, Maraun, Scheu, Rillig, & Caruso, 2015; Nielsen et al., 2010). Besides, the soil environment limits mobility and sensing of chemical cues by predators / consumers, which complicates selective feeding on specific food resources. ‘Convergence‘ in Δ^13^C values in different taxa suggests that different types of organic compounds (plant tissues or microbially processed detritus; Potapov et al., 2018) are utilized by an array of different lineages of consumers resulting in functional redundancy. Resource specialization such as feeding on different organic matter compounds, therefore is unlikely to drive evolutionary adaptations in soil invertebrates, suggesting that it is relatively easy to switch between herbivory and detritivory or between preying on herbivores and decomposers. Such switches have been shown e.g., for species of Collembola (Endlweber, Ruess, & Scheu, 2009), Chilopoda (Klarner et al., 2017) and Elateridae (Samoylova & Tiunov, 2017).

The stronger taxonomic signal in Δ^15^N as compared to Δ^13^C values suggests that switching between prey of different trophic levels is evolutionary difficult resulting in taxa being conserved within their trophic level. Despite there are examples of predatory species in detritivore lineages such as Collembola (Hopkin, 1997; Potapov et al., 2016) and Oribatida (Heidemann et al., 2014; Maraun et al., 2011), they likely represent only a minority of the species of these groups. Notably, isotopic dissimilarity increased steadily from the species to the class level, but decreased thereafter, suggesting that the differences in trophic level between taxa were established early in the evolution of these lineages, i.e. during the colonisation of land by the major arthropod lineages, and diversification within these lineages was associated by refinement of trophic niches (Rota-Stabelli et al., 2013; Schaefer, Norton, Scheu, & Maraun, 2010).

### Taxonomic sufficiency in soil food-web studies

Species is a keystone but one of the most debated concepts in biology. For instance, the increasing recognition of cryptic species challenges morphology as the criterion for species delineation (Fišer, Robinson, & Malard, 2018). However, virtually all animal and plant taxa are described based on morphology (Cook, Edwards, Crisp, & Hardy, 2010). Taxonomists delineate supraspecific taxa from each other in order to establish a system of morphologically consistent units. Normal and unimodal distributions of isotopic niches of genera and even families, suggest that the taxonomic system developed for soil invertebrates is consistent also in ecological functions and thus may be used in ecological studies. This, however, needs to be done with caution since each question addressed may deserve its own level of taxonomic resolution (Bhusal, Kallimanis, Tsiafouli, & Sgardelis, 2014; Timms et al., 2013). To facilitate selection of the adequate taxonomic level for respective research questions we provide a list of mean Δ^13^C and Δ^15^N values of soil invertebrates from temperate forests from species to class level of taxonomic resolution and classified the records according to their reliability and trophic flexibility in the Appendix (Tables S1-S4).

Among taxa, maximum information on the structure of trophic niches is obtained at the species level of taxonomic resolution. Nevertheless, our data suggest that species identity in soil communities provides little additional information relative to the respective genera. In 92% of cases stable isotope composition of congeneric species was not significantly different and the mean offset of populations within genera was only 0.1‰ higher than that of populations within species. For instance, trophic niches in congeneric species of Chilopoda and Lumbricidae were shown to be similar (Ferlian, Scheu, & Pollierer, 2012; Schmidt, Curry, Dyckmans, Rota, & Scrimgeour, 2004). Also, congeneric species of Collembola and Oribatida typically have similar stable isotope composition, although there are exceptions (Potapov et al., 2016; Schneider et al., 2004). The cases in which the stable isotope composition of congeneric species differed significantly (8%) were distributed across a variety of taxa including Oribatida, Collembola and Formicidae (see Table S2). Niche differentiation among congeneric species may in part be explained by morphological variability within genera, pointing to the need to disclose relationships between morphological traits and feeding habits of soil invertebrates in future studies (Malcicka, Berg, & Ellers, 2017; Potapov et al., 2016).

The rather small loss of information on trophic niches when using genera instead of species in large was due to the high variability in stable isotope composition among populations within species, which was emphasized before (Lehmitz & Maraun, 2016; Tillberg, McCarthy, Dolezal, & Suarez, 2006; Zalewski et al., 2014). A recent study based on molecular gut content analysis of three morphologically similar species of Collembola showed high diversity and temporal variability in their fungal diet, but no significant differences in feeding habits were found between species across locations and seasons (Anslan, Bahram, & Tedersoo, 2018). Clearly, the trophic niche of species / taxa in a given community will always depend on the biotic and abiotic environment and it is preferable to explicitly study the trophic niches of species of the community investigated.

Δ^13^C and Δ^15^N values of species within families were not different in 80 - 60% of the cases indicating that trophic niches of species are not well represented using families as functional units as suggested e.g., for Mesostigmata in grasslands (Walter & Ikonen, 1989). The trophic differentiation of species / taxa showed different trends in the three major lineages of soil arthropods. Chelicerata (represented by Arachnida) and Myriapoda were enriched in ^13^C and ^15^N as compared to Hexapoda, with the enrichment being on average 0.8‰ for both Δ^13^C and Δ^15^N values. However, large variations in ^13^C and ^15^N values suggest large variability in trophic niches in each of the three lineages. Myriapoda in our dataset comprised two classes, i.e. Diplopoda and Chilopoda feeding on very different resources, i.e. dead organic matter and animal prey, respectively. Arachnida in our dataset comprised Oribatida, predominantly living as detritivores and fungal feeders, and Mesostigmata and Araneae living as predators. In Hexapoda the trophic niche varied between classes (Insecta vs. Collembola) and also between orders. Presumably, this is related to the high diversity of Hexapoda exceeding that of Arachnida and Myriapoda (Zhang, 2011). Notably, the Δ^13^C and Δ^15^N values of species-rich taxa such as Coleoptera varied markedly due to including taxa of very different trophic position such as Carabidae living mainly as predators and Tenebrionidae living mainly as detritivores (Table S1). This supports the notion that species- and genus-rich families need to be resolved at higher taxonomic resolution to adequately represent their trophic position (Jiang et al., 2013; Timms et al., 2013). A wider range of niches in species-rich taxa illustrates a close link between morphological and ecological diversification. The diversification of lineages is driven primarily by the occupation of new niche space (Gavrilets & Losos, 2009; Mahler, Revell, Glor, & Losos, 2010). During the long evolutionary history of old lineages, some lineages, such as Insecta, managed to exploit novel resources / prey species thereby radiating markedly, whereas other lineages, such as Diplopoda and Chilopoda, remained confined to their trophic niche and remained less species rich. Therefore, using supraspecific taxa as functional units in ecological studies needs to consider the degree of ecological diversification of the different lineages rather than using a uniform taxonomic level across the taxa studied. Including Δ^13^C and Δ^15^N values as continuous variables allows to refine classic food-web models based on discrete trophic levels and to include omnivory and intraguild predation which are of significant importance for food web dynamics in soil (Digel et al., 2014).

## Conclusions

Soil animals that are sharing similar taxonomic affiliation and therefore morphology are also sharing similar trophic niches. Blomberg’s K criteria suggest that feeding on different basal resources (plants or detritus) is ‘convergent’ across different lineages of soil invertebrates whereas the occupation of different trophic levels followed the Brownian-motion taxonomic model. We found no evidence for strict trophic niche conservatism in soil invertebrates.

Although higher taxonomic resolution provides more information on feeding habits, in 92% of the cases stable isotope values of congeneric species of soil invertebrates did not differ significantly. This small loss of information was due to the high variability in stable isotope composition between populations within species. Nevertheless, trophic niches of populations within genera and families fitted normal and unimodal distributions, suggesting trophic consistency of these supraspecific taxa. Supraspecific taxa can serve as meaningful functional units in ecological studies, but the choice of taxonomic resolution needs to be adopted based on the particular research question and taxonomic group. In particular, diverse groups, such as Oribatida, Collembola and Coleoptera, deserve higher taxonomic resolution. Further studies need to explore how and why trophic radiation varies between lineages. The compiled list of mean Δ^13^C and Δ^15^N values of taxa can be combined with other literature data in order to infer feeding habits of soil invertebrates and move towards more realistic soil food-web models.

## Acknowledgements

The study was supported by the Russian Foundation for Basic Research, Project #18-04-01200 and in part by the Deutsche Forschungsgemeinschaft (DFG) in the framework of the Collaborative Research Centre CRC 990. We thank Ting-Wen Chen for advices on data analysis.

## Author contributions statement

AP analysed the data and worked on the drafts. AP, AT and SS conceived and refined the ideas. All authors contributed critically to the drafts and gave final approval for publication.

## Data accessibility statement

Should the manuscript be accepted, the data supporting the results will be archived in a public repository and the data DOI will be included at the end of the article.

